# Pheophorbide *a*, a chlorophyll catabolite may regulate jasmonate signalling during dark-induced senescence in *Arabidopsis*

**DOI:** 10.1101/486886

**Authors:** Sylvain Aubry, Niklaus Fankhauser, Serguei Ovinnikov, Krzysztof Zienkiewicz, Ivo Feussner, Stefan Hörtensteiner

## Abstract

Chlorophyll degradation is one of the most visible landmarks of leaf senescence. During senescence, chlorophyll is degraded in the multi-step pheophorbide *a* oxygenase (PAO)/phyllobilin pathway, which is tightly regulated at the transcriptional level. This regulation allows a coordinated and efficient remobilisation of nitrogen towards sink organs. Taking advantage of combined transcriptome and metabolite analyses during dark-induced senescence of *Arabidopsis thaliana* mutants deficient in key steps of the PAO/phyllobilin pathway, we show an unanticipated role for one of the pathway intermediates, *i.e*. pheophorbide a. Both jasmonic acid-related gene expression and jasmonic acid precursors specifically accumulated in *pao1*, deficient in PAO. We propose that pheophorbide a, the last intact porphyrin intermediate of chlorophyll degradation and unique pathway ‘bottleneck’, has been recruited as a signalling molecule of the chloroplast metabolic status. Our work challenges the assumption that chlorophyll breakdown is merely a senescence output, but propose that the flux of pheophorbide *a* through the pathway acts in a feed-forward loop that remodels the nuclear transcriptome and controls the pace of chlorophyll degradation in senescing leaves.

**Summary:** Transcriptome and metabolite profiles of key chlorophyll breakdown mutants reveal complex interplay between speed of chlorophyll degradation and jasmonic acid signalling

**Financial sources:** This work was supported by the European Union (Plant Fellow program), the Swiss National Foundation/ERA-NET (grant N° 163504) and the German Research Foundation (DFG, grant N° INST 186/822-1).

## INTRODUCTION

In higher plants, leaf senescence is a tightly regulated process that is responsible for remobilisation of nutrients like nitrogen and phosphorus from source to sink organs (Hörtensteiner and Feller, 2002). Degradation of photosynthetic proteins, representing up to 70% of total leaf proteins, is co-regulated with chlorophyll (chl) degradation, while carotenoids are largely retained (Kusaba et al., 2009). Chl is degraded via a cascade of coordinated enzymes leading to cleavage and export of chl catabolites to the vacuole in the form of non-toxic linear tetrapyrroles, termed phyllobilins (Süssenbacher et al., 2014). Since all final phyllobilins are ultimately derived from the porphyrin ring-opening activity of PHEOPHORBIDE A OXYGENASE (PAO), this pathway of chl breakdown is referred to as the “PAO/phyllobilin pathway” (Hörtensteiner, 2006).

Two key chlorophyll catabolic genes (CCGs) that encode the chlorophyll catabolic enzymes (CCEs) and precede the opening of the porphyrin ring of chl, *i.e*. the magnesium dechelating enzyme NON YELLOWING (NYE) and PHEOPHYTIN PHEOPHORBIDE HYDROLASE (PPH), hydrolyzing the phytol tail, are tightly co-regulated with PAO at the transcriptional level during leaf senescence. In addition, all three CCEs were shown to physically interact (Pružinská et al., 2007; Ren et al., 2007; Aubry et al., 2008; Sakuraba et al., 2012). This regulation may allow quick metabolic channelling of potentially phototoxic chl catabolites. A model based on the apparent coordinated expression of these genes and under the control of one or a few main transcriptional regulator(s) could therefore be hypothesised. However, the mechanism underlying this transcriptional coordination remains unclear.

Senescence is a complex process integrating hormonal and environmental signals from very distinct pathways (Kim et al., 2018). Only considering CCGs, at least three distinct hormonal signals and their respective signalling pathways have been shown to interact via some of their components with CCG promoters (for a recent review, see (Kuai et al., 2018))(Kuai *et al.*, 2017)(Kuai *et al.*, 2017). Jasmonic acid (JA), ethylene (ET) and abscisic acid (ABA) signalling pathways together with some components of the light signalling cascade have been shown to modulate CCG expression directly (Kuai et al., 2018).

In particular, JA and its derivatives are key regulators of senescence (He *et al.*, 2002) and typically synthesised in response to insects and necrotrophic pathogens (Kim et al., 2018; Wasternack and Feussner, 2018). Levels of JA increase during natural or dark-induced senescence (Breeze et al., 2011) and ectopic methyl jasmonic acid induces early senescence (Ueda and Kato, 1980). JA and associated oxylipin signalling have pleiotropic effects on the cellular fate, for example changing expression of defense genes (Hickman et al., 2017). Default JA signalling is perceived via CORONATINE-INSENSITIVE 1 (COI1) that in turn degrades the transcriptional repressors JA ZIM-domain proteins (JAZ) (Howe et al., 2018). For example, JAZ7 blocks MYC2 transcription factor activity that act upstream of many genes involved in dark-induced leaf senescence (Yu et al., 2016). MYC2/3/4 and their downstream targets NAC019/055/072 directly interact with NYE1, NYE2 and NYC1 promoters (Zhu et al., 2015). In a very similar manner, NAC019 and MYC2 interact with each other to synergistically up-regulate NYE1 (Zhu et al., 2015). Other transcription factors involved in ET signalling (EIN3, EEL, ORE1 and ERF17), ABA signalling (NAC016, NAC046, NAP, ABF2/3/4, ABI5) were also reported as direct interacting transcription factors of some *cis*-elements in CCG promoters (Kim et al., 2014; Sakuraba et al., 2014; Qiu et al., 2015; Sakuraba et al., 2016; Yin et al., 2016). These multiple intertwined hormonal cues eventually lead to chlorosis, by way of degradation of chl, as a visible landmark of dark-induced, aged-induced and also (a)biotic stress-induced senescence. Interestingly, constitutive overexpression of single CCGs in *Arabidopsis thaliana* (Arabidopsis) led in most cases to an acceleration of chl breakdown after senescence induction (Sakuraba et al., 2012). This suggests a feedback mechanism by which the chloroplast coordinates the rate of chl degradation during leaf senescence. The extent to which the speed of chl degradation itself could regulate the various hormonal cues and thereby inform cells about the current status of their senescing chloroplasts remains to be shown.

Here, in an attempt to identify such a link and simultaneously shed more light onto these complex regulatory networks, we used genome-wide transcriptome analysis of CCG mutants during early dark-induced senescence. By combining these data with metabolite profiling, we aim at understanding processes that regulate the dynamics of the production of chl catabolites in the PAO/phyllobilin pathway and the extent to which accumulation of pathway intermediates remodel nuclear gene expression, and more precisely the JA response.

Based on our data, we propose a model where transient accumulation of the intermediate pheophorbide (pheide) *a* acts as a sensor for the rate of chl degradation, and thereby regulates the speed of leaf senescence tuned by JA signalling. This model highlights a new function for the PAO/phyllobilin pathway of chl breakdown, not only as an irreversible prerequisite to senescence-driven nitrogen remobilization, but also as a sensing mechanism of the stress status of the chloroplast.

## RESULTS

### Coordinated Variations of the Leaf Transcriptome During Dark-Induced Senescence

The extent of variations in gene expression during dark incubation of detached leaves (DET) was assessed using RNAseq. Mature leaves number six (see Materials and Methods) were sampled in triplicate at 0 and 2 days in the dark (dd). Using these time points allowed us to profile early events of the senescence program before any distinct visible phenotype (Fig. 1A).

**Figure 1.**
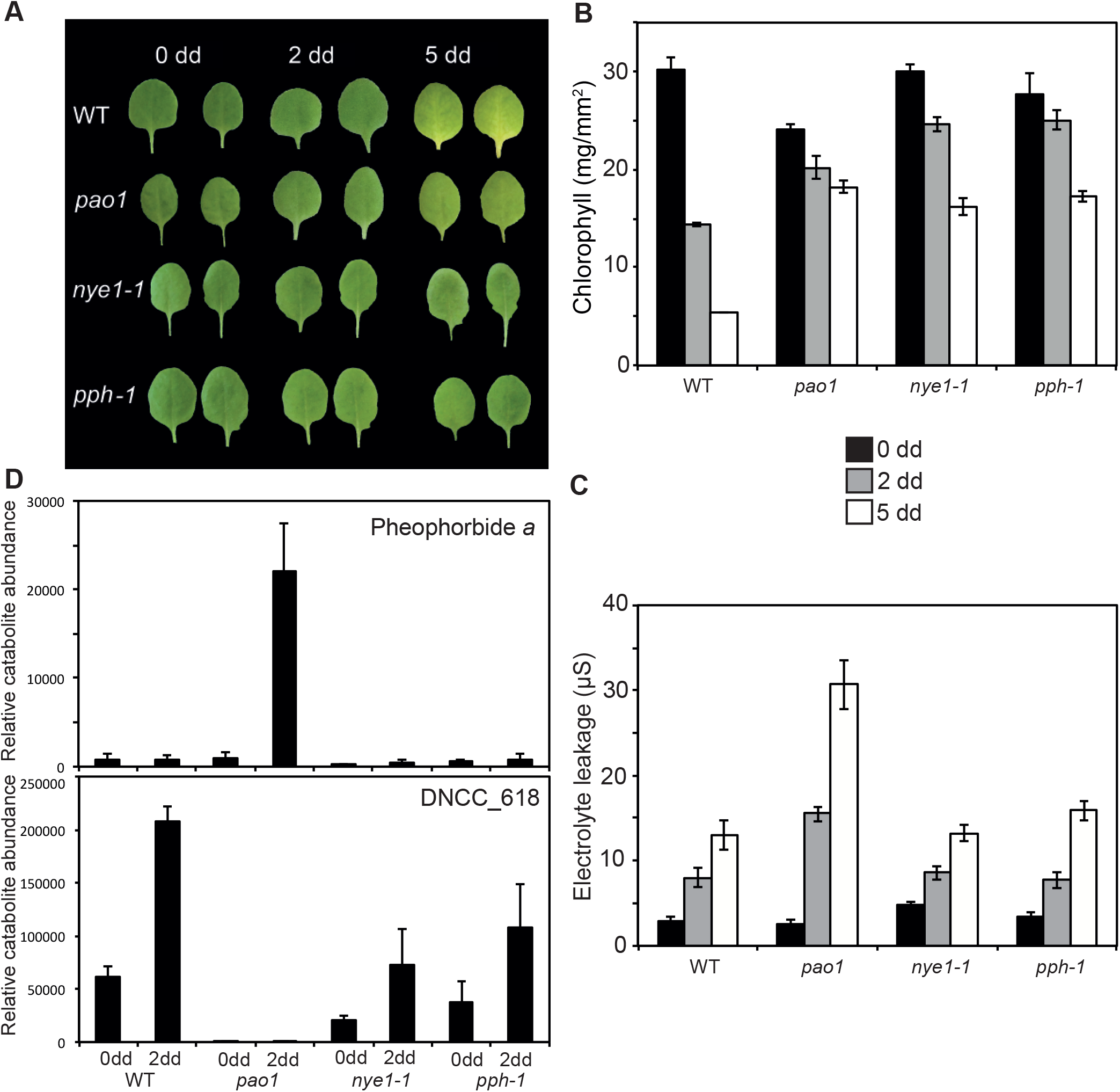
Phenotypic characterisation of CCGs mutants during dark-induced senescence of detached leaves. **A** WT, *pao1*, *nye1-1* and *pph-1* detached leaves before and after 2 and 5 days of dark induced senescence (dd). **B** Chlorophyll degradation of CCG mutants during dark-induced senescence. **C** Electrolyte leakage of CCG mutants during dark-induced senescence. **D** Profile of the accumulation of pheophorbide *a* and the major phyllobilin (DNCC_618) in CCG mutants during dark-induced senescence. Data in **B-D** are mean values of a representative experiment with three (**B**), at least ten (**C**) and five (**D**) replicates, respectively. Error bars indicate SD.

In wild type (WT) leaf, a total of 21,403 genes were detected (genes with normalised counts ≥ 1 in at least one of the samples), amongst these 6,124 (29% of detected) genes were considered as being differentially expressed during DET (after applying EBSeq test using a posterior probability of differential expression ≥ 0.95 and a minimum fold change of two times). Of these, 3,389 genes were upregulated after dark treatment (Table 1). In an analogous experiment on Arabidopsis leaf senescence using microarrays, 2,153 genes were differentially expressed between 0 and 2 dd, of which 65% (1,353) were common to our dataset (Supplemental Fig S1) (Van der Graaff et al., 2006). Another microarray-based analysis of the transcriptome during natural leaf senescence in Arabidopsis showed perturbation in 6,370 genes, of which 2,825 (44%) were common to our differentially expressed genes Supplemental Fig S1) (Breeze et al., 2011). These results show the biological relevance of our data. The differences observed being likely due to the biases associated with different techniques used to induce senescence. A relatively high number of genes have been shown to be similarly expressed when comparing different methods of senescence induction, such as DET, dark incubation of attached leaves and natural senescence (Van der Graaff et al., 2006). Analysis of the WT transcriptome signature using Gene Ontology (GO) terms revealed that photosynthesis, starch metabolism, glucosinolates, tetrapyrrole synthesis and redox terms were under-represented during DET (Fig. 2A), while terms gathering genes involved in lipid, amino acid and protein degradation but, more interestingly, also micro-RNA, retrotransposons and the bZIP family of transcription factors were over-represented during senescence (Fig. 2A). This is consistent with described major gene expression changes during leaf senescence (Van der Graaff et al., 2006; Breeze et al., 2011).

**Table 1.** Number of genes differentially expressed during dark incubation of detached leaves (DET) in WT and three CCG mutant lines. 25,920 genes were detected in at least one of the 24 samples.

| Total transcripts detected (non zero) | 21,403 |  |  |
| --- | --- | --- | --- |
| Differentially expressed during DET<br>(PPDE $\geq 0.95$ and FC $\geq 2$) | Total | Up-regulated | Down-regulated |
| WT | 6,124 | 3,389 | 2,735 |
| *nye1-1* | 6,227 | 3,325 | 2,902 |
| *pph-1* | 6,764 | 3,777 | 2,987 |
| *pao1* | 11,408 | 5,723 | 5,685 |

**Figure 2.**
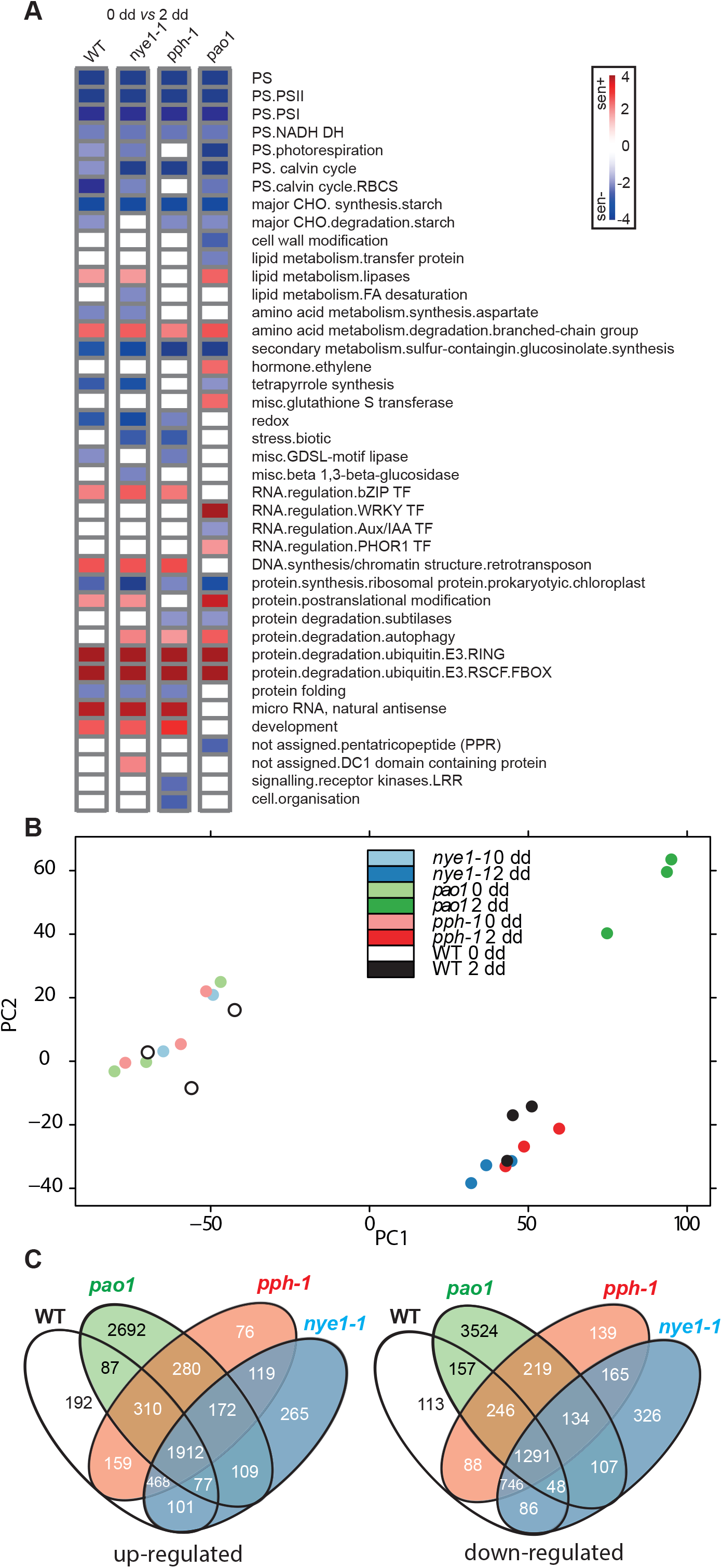
RNAseq profiling of CCG mutants provide new insight into the relationship of the PAO/phyllobilin pathway to global leaf senescence. **A** Major enriched gene ontology terms identified in the three CCG mutants during dark-induced senescence (0 dd vs 2 dd) using Wilcoxon test implemented in Pageman tool (Usadel et al., 2006). **B** Principal Component Analysis of the RNAseq data. **C** Venn diagrams showing common patterns of differential expression (0 dd vs 2 dd) of up- and down-regulated genes during dark-induced senescence.

### Disruption of Specific CCGs Modifies Phyllobilin Accumulation that Lead to Distinct Stay-Green Phenotypes

In order to determine the extent to which the disruption of the PAO/phyllobilin pathway influences the global leaf senescence process, we analysed three mutants that are defective in this pathway: *nye1-1*, *pph-1* and *pao1*, and analysed senescence of detached leaves after dark incubation (Pružinská et al., 2003; Ren et al., 2007; Schelbert et al., 2009). After 2 dd, chl was retained in these lines (Fig. 1A and B). Noteworthy, *pao1* contained slightly less chl before dark incubation (0 dd) in comparison to WT (Fig. 1B). In addition to chl retention, *pao1* accumulated pheide *a* during dark incubation and exhibited a light-independent cell death (LICD) phenotype (Pružinská et al., 2003; Hirashima et al., 2009), as deduced from an increase in electrolyte leakage of *pao1* leaf tissue in the dark (Fig. 1C). The molecular basis of LICD in this line is unclear, but this phenotype is specific to *pao1*, and may to some extent be linked to pheide *a* accumulation (Fig. 1D) (Hirashima et al., 2009). Absence of PAO in *pao1* led to a complete halt of the PAO/phyllobilin pathway with virtually no phyllobillin accumulation (Fig. 1D). By contrast, *nye1-1* and *pph-1* accumulated the major phyllobilins of Arabidopsis to about one third of the WT level after two days of dark treatment but virtually no pheide *a* (Fig. 1D) (Christ et al., 2013).

Stopping the PAO/phyllobilin pathway artificially at various levels seems to imply largely distinct phenotypic modifications. On the top of its relatively well described implication on nitrogen remobilisation (mostly due to photosystem degradation), the control of chl catabolite homeostasis within the degradation pathway is a potentially overlooked signal that may inform the cell about the status of chloroplast integrity or metabolism.

### Transcriptome Analysis of CCG Mutants Gives Insight Into Molecular Bases of Phenotypic Variations Observed in the Dark

We then assessed alterations in the leaf transcriptome during DET in all three mutants. In *nye1-1* and *pph-1*, 6,227 and 6,764 genes were differentially expressed between 0 and 2 dd, respectively, numbers that were comparable to the changes observed in WT (Table 1). By contrast, about two times more genes (11,408) were differentially expressed in *pao1* during DET (Table 1). In order to detect variations in gene expression that were specific to mutations of one or several of the CCGs, genes differentially expressed during DET in every line were compared (Fig. 2B, C). 2,692 and 3,524 genes, respectively, were specifically down- and up-regulated in *pao1*, while for *pph-1* (76 and 139 genes, respectively) and *nye1-1* (265 and 326 genes, respectively) these numbers were much smaller (Fig. 2C). A core set of 3,203 genes (1,912 up- and 1,291 down-regulated) showed similar patterns of expression in all four lines. The most enriched genes among these were genes involved in catabolic processes, senescence, aging and autophagy, while genes involved in chloroplast and various photosynthesis-related processes were the most repressed ones (Supplemental Dataset 2). Analysis of GO term enrichment showed that altered genes showing a very similar pattern in all four lines included genes involved in photosynthesis, starch metabolism, glucosinolate synthesis (down-regulated) as well as protein and amino acid degradation (up-regulated) (Fig. 2A). Collectively, this indicated that mutations in any of the three CCGs, despite clear phenotypic differences in these lines, had little effect on general background senescence processes.

Most of the genes whose expression specifically changed in *pao1* while remaining unchanged in all other lines belonged to GO terms related to ethylene and WRKY and PHOR1 transcriptional regulators (Fig. 2A). Among GO terms that were significantly enriched in *pao1*, categories of genes involved in various stresses were the most enriched ones: these include response to stress, stimulus, chitin, carbohydrate, chemical stimuli as well as genes involved in post-transcriptional processes (Supplemental Dataset 2).

Thirty-six of the 50 most highly expressed genes after 2 dd were different between *pao1* and WT (Supplemental Dataset 3), among them, PLEIOTROPIC DRUG RESISTANCE 12 (PDR12/ABCG40), involved in ABA transport (Kang et al., 2010)(Kang *et al.*, 2010)(Kang *et al.*, 2010), LIPOXYGENASE 1 (LOX2) involved in JA synthesis (Wasternack and Feussner, 2018), as well as NYE1. Taken together our data suggest major remodelling of gene expression in *pao1* leaves upon dark incubation, while absence of NYE1 or PPH only mildly affect the senescence leaf transcriptome, at least at an early stage of senescence.

### The PAO/Phyllobilin Pathway Is Mainly Regulated at the Transcriptional Level

Most of the core CCGs like PAO, *PPH* and *NYE1*, as well as genes encoding some catabolite-modifying enzymes, *i.e*. METHYLESTERASE 16 (MES16) and CYTOCHROME P450 MONOOXYGENASE 89A9 (CYP89A9) were transcriptionally up-regulated during DET in WT (Supplemental Dataset 1 and Fig. 3) (Sakuraba et al., 2012). In all four lines studied, genes encoding enzymes involved in the oxidative half of the chl cycle, namely CHLOROPHYLL *A* OXYGENASE (CAO) and CHLOROPHYLL SYNTHASE (CHLG), were down-regulated, whereas genes involved in chl *b* to chl *a* conversion (*NYC1* and NOL) were up-regulated. This is consistent with the assumption that conversion of chl *b* to chl *a* is a prerequisite for chl degradation (Sakuraba et al., 2010). Noteworthy, expression of *RCCR* was repressed during DET and, thus, not correlated with the expression of *PAO* or of any of its proposed interacting partners (Fig. 3) (Sakuraba et al., 2012). Except for a slight decrease in *HCAR*, *nye1-1* and *pph-1* did not exhibit significant differences in CCG expression compared to WT. By contrast, major changes were observed in *pao1* with strong overexpression of *NYE1* and *NYC1* (but not *NOL*) and down-regulation of *CAO* and *HCAR*, suggesting a “feed-forward” regulation of the catabolic pathway.

**Figure 3.**
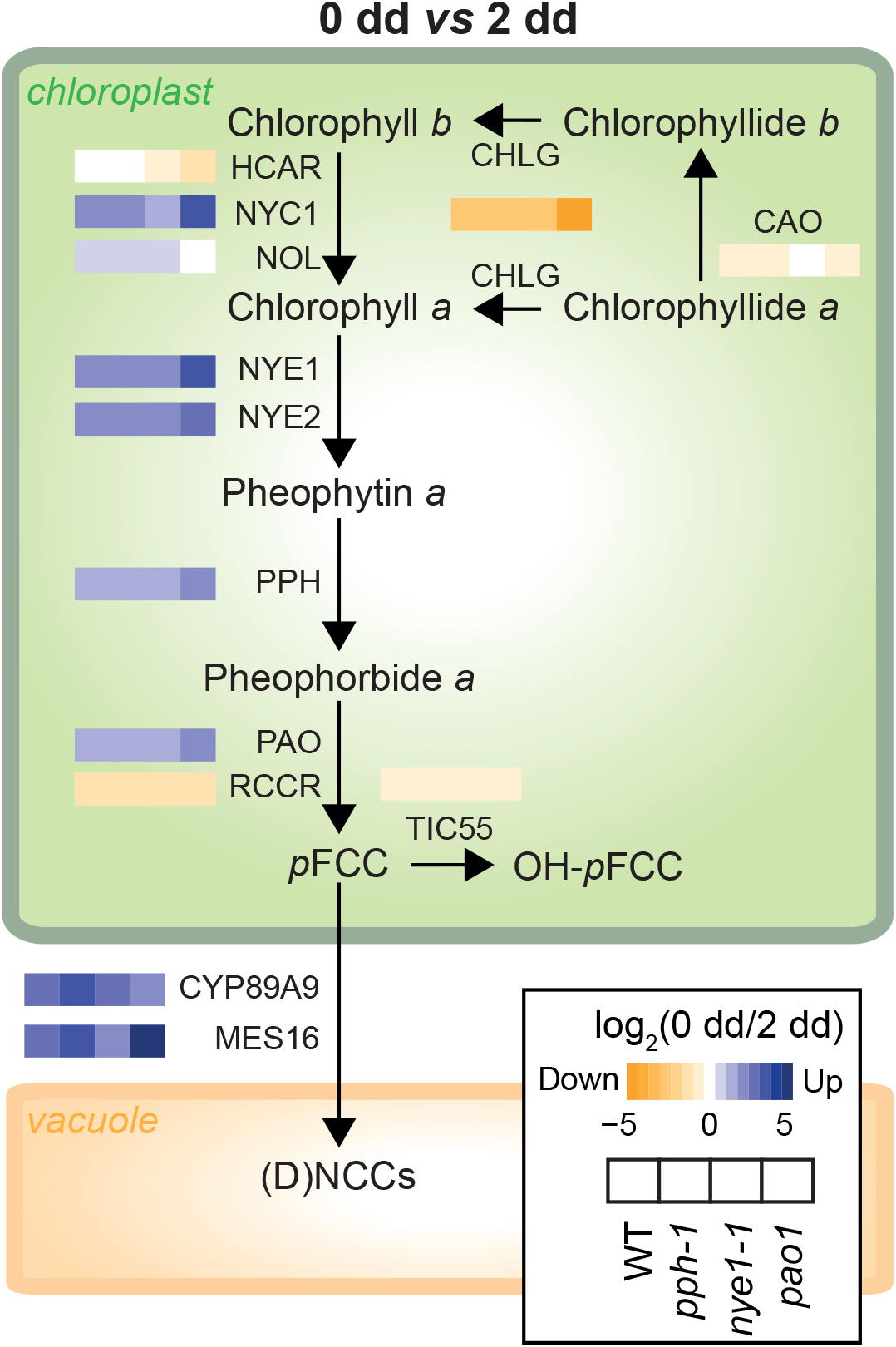
Influence of dark-induced senscence on the expression of the genes involved in the PAO/phyllobilin pathway. Heat maps represent log_2_ (fold change) of gene expression in each of the four studied lines during dark-induced senescence. Genes/enzymes: CAO, chlorophyll *a* oxygenase; CHLG, chlorophyll synthase; CYP89A9, cytochrome P450 monooxygenase 89A9; HCAR, 7-hydroxymethyl chlorophyll *a* reductase; MES16, methylesterase 16; NYC1, non-yellow coloring 1 (chlorophyll *b* reductase); NYE1, non yellowing 1 (magnesium dechelatase); NYE2, non yellowing 2; PAO, pheophorbide *a* oxygenase; PPH, pheophytinase; RCCR, RCC reductase; TIC55, translocon at the inner chloroplast envelope 55. Phyllobilins: DNCC, dioxobilin-type NCC; NCC, non-fluorescent chlorophyll catabolite; pFCC, *primary* fluorescent chlorophyll catabolite; RCC, red chlorophyll catabolite.

### The PAO/Phyllobilin Pathway Is Controlled by Multiple Intertwined Signalling Pathways

In order to evaluate the impact of CCG mutations on upstream regulators of the pathway, we extracted expression data for signalling pathways involving JA, ET, ABA and light signalling in all four lines as described (Kuai et al., 2018) (Fig. 4). Out of the 41 genes represented here that are key genes involved in these hormonal pathways, only *JAZ10* was significantly downregulated in *pao1* as compared to WT (none in *pph-1* or *nye1-1*, Supplemental Dataset 5), suggesting minor function of the hormonal cues before dark incubation. Expression of genes involved in ET and ABA signalling was mostly up-regulated in all lines during dark-induced senescence. The most striking difference between *pao1* and all other three lines was the pattern of expression of genes involved in JA signalling: *COI1* expression was increased significantly during dark treatment in WT, *pph-1* and *nye1-1*, but not in *pao1*, while nine of the twelve jasmonate-ZIM domain (JAZ) proteins showed an inverse pattern of expression. Intriguingly, among the very few genes differentially expressed after dark treatment in both *nye1-1* and *pph-1*, a subset of JAZ genes, namely *JAZ1*, *JAZ5*, *JAZ7*, *JAZ8* and *JAZ10*, were significantly down-regulated compared to WT (Supplemental Dataset 1).

**Figure 4.**
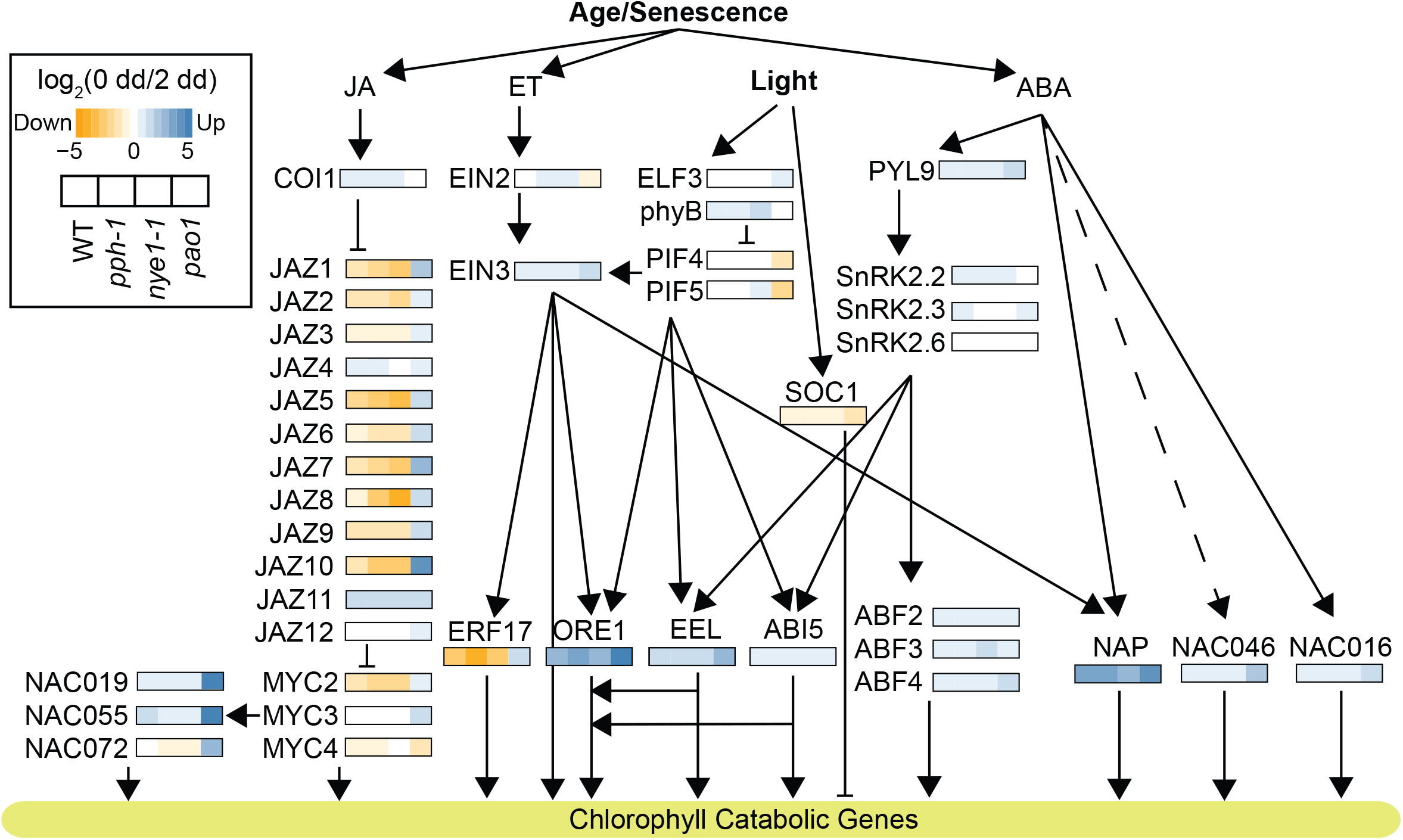
Transcriptional regulation of the PAO/phyllobilin pathway during dark-induced senescence is mainly affected in *pao1*. Heat maps represent log_2_ (fold change) of gene expression in each of the four studied lines during dark-induced senescence. JA, jasmonic acid; ET, ethylene; ABA, abscisic acid; COI1, coronatine insensitive 1; JAZ, jasmonate-ZIM domain; NAC, NAM, ATAF1/2 and CUC2 domain protein; NAP, NAC-like, activated by PA3/PI; EIN, ethylene insensitive; ELF3, early flowering 3; PIF, phytochrome interacting factor; SOC1, suppressor of overexpression of coi1; ERF17, ethylene response factor; ORE1, oresara 1; EEL, enhance em level; ABI5, ABA insensitive 5; ABF, ABA-responsive element binding factor; SnRK2, serine/threonine kinase 2; PYL9, pyrabactin resistance 1-like 9.

It also appears that expression of transcription factors that are repressed by JAZ proteins like MYC2/3 and downstream factors like NAC019/055 and 072 were also up-regulated exclusively in *pao1* at 2 dd (Fig. 4).

### Accumulation of Pheide *a* in *pao1* Modifies JA-Related Signalling

Having noticed strong variations of JA-related gene expression in *pao1* after dark incubation, we analysed whether JA synthesis and levels of JA metabolites were also modified in this line. To this end, JA precursors (12-OPDA, dn-OPDA, OPC6, and OPC4) as well as JA and some of its derivatives (JA-Val, JA-Ile, JA-Leu, 12OH-JA-Ile, 12COOH-JA-Ile, 12O-Glc-JA and 12HSO_4_-JA) were quantified in both WT and *pao1* (Fig. 5 and Supplemental Dataset 4). JA levels were significantly increased in WT during dark incubation, but in *pao1*, JA accumulated with an order of magnitude higher, *i.e*. up to 2 nmol g^-1^ fresh weight (Fig. 5 and Supplemental Dataset 4). Levels of endogenous JA after dark treatment are known to increase in WT (Seltmann et al., 2010a) and are regulated under strong circadian control (Goodspeed et al., 2012). However, the dark-induced increase of JA in WT not necessarily triggers JA-signalling pathways (Seltmann et al., 2010b). In *pao1*, not only JA levels were dramatically increased, but also downstream metabolites, *i.e*. JA-Val, JA-Leu, 12OH-JA-Ile and the active phytohormone JA-Ile (Fig. 5 and Supplemental Dataset 4). Genes involved JA biosynthesis (*LOX2*, *AOC1*, *AOC2*, *OPR3)* and degradation (*CYP94B1*, *CYP94B3*) were also strongly upregulated in *pao1*. Interestingly, expression of *JMT* and *JAR1* were unchanged in both lines.

**Figure 5.**
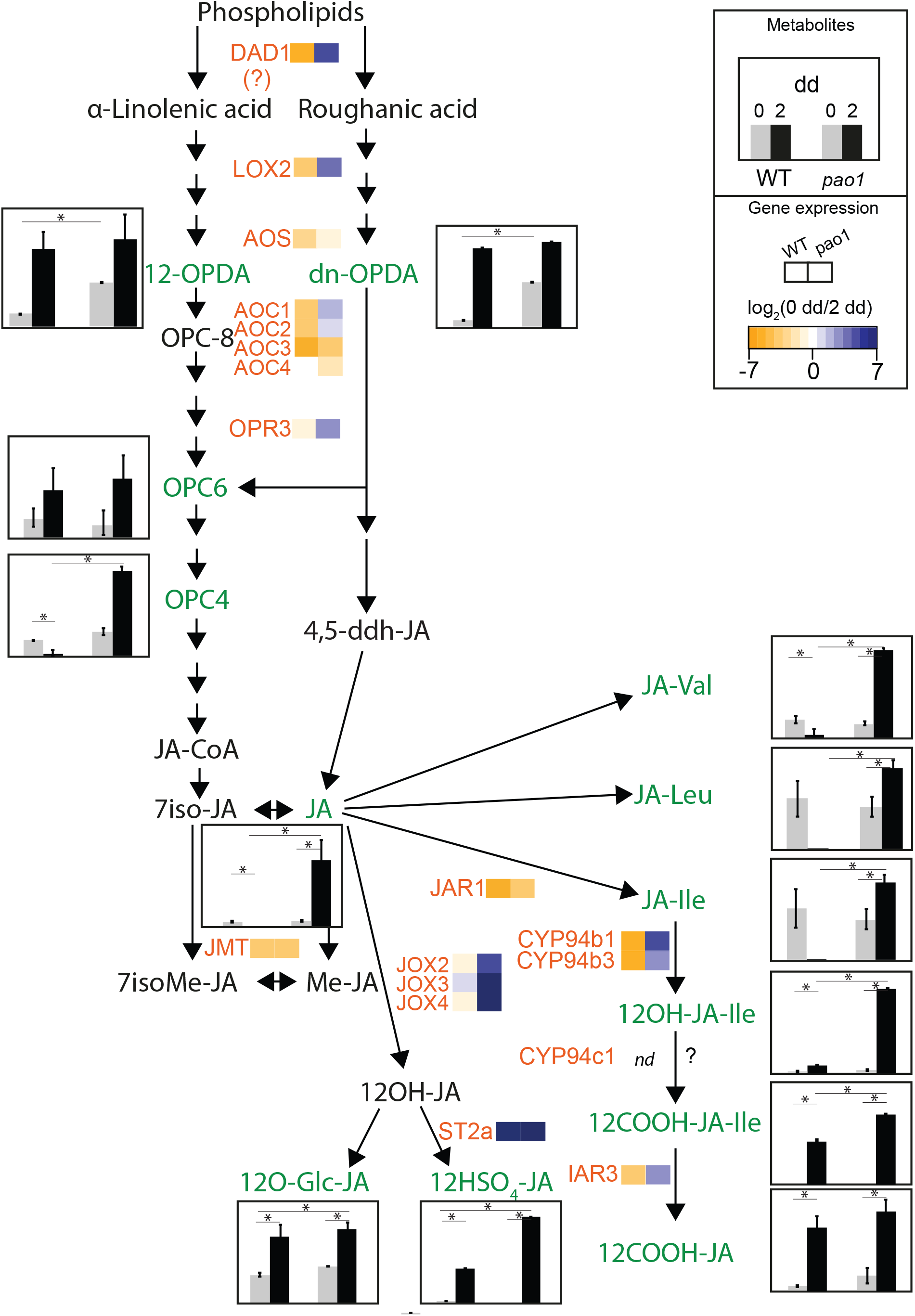
Jasmonic acid metabolism during dark-induced senescence in WT and *pao1*. Levels of JA and JA-related metabolites in grey (0 dd) and black (2 dd) for WT and *pao1* are shown as histograms. Expression levels are shown using heat maps of log_2_(fold change). Genes/enzymes: DAD1, delayed anther dehiscence 1; LOX2, 13-lipoxygenase 2; AOS, allene oxide synthase; AOC, allene oxide cyclase; OPR3, OPDA reductase 3; IAR3, IAA-alanine resistant 3; JMT, jasmonate methyltransferase; JAR1, JA-amino acid synthetase 1; JOX, JA-induced oxygenase; CYP, cytochrome P450 monooxygenase; ST2A, sulfotransferase 2. Metabolites: OPDA, 12-oxo-phytodienoic acid; OPC 3-oxo-2-cis-2-pentenyl cyclopentyl-octanoic acid; JA-CoA, jasmonate-coenzyme A. Asterisks indicate significant differences (p < 0.05).

Taken together, these data show a complex rewiring of the JA signalling pathway and indicate a link between the *pao1* phenotype and JA responses. Next, we tried to decipher the exact extend of this feedback using patterns of gene co-expression.

### Co-Expression Analysis Reveals Structure of Regulatory Networks of the PAO/Phyllobilin Pathway

To further characterise a possible link between the PAO/phyllobilin pathway and JA signalling, we computed the genome-wide expression data for all four lines studied and tried to decipher co-expression patterns underpinning relevant gene networks. The basic assumption being that genes that show a similar pattern of expression during DET and/or in various genetic backgrounds could be involved in a similar process and most probably share similar regulating pathways. We used Weighted Genome Co-expression Network Analysis (WGCNA) (Zhao et al., 2010) to perform comparative analysis of gene co-expressed modules among darkness treatment in all four lines. Genes were clustered in 16 co-expression modules, each harbouring genes that generally showed a similar pattern of expression across genetic background and treatment (Fig. 6A, Supplemental Fig. S2). Three modules (blue, pink and yellow) were highly correlated with the darkness treatment. The pink and yellow modules contained genes that showed consistent changes expression during dark incubation in all four lines, but not the blue motif that did not correlate to *pao1* after 2 dd (Fig. 6B). Modules were subsequently characterised using GO term enrichment (Fig. 6B & Supplemental Dataset 6). All three motives were enriched in terms related to mRNA catabolic process, fatty acid catabolism, senescence and autophagy (Supplemental Dataset 6). Red, black and green modules that mostly correlated with *pao1* after dark incubation were enriched in terms representing various responses to stress as well as hormonal response (namely ET, JA and ABA responses, Supplemental Dataset 6). Interestingly, *PPH* is the hub gene. *i.e*. the most highly connected gene in this module (Langfelder and Horvath, 2008), of the blue module that contains most CCGs (*PAO*, *CYP89A9*, *PPH*, *NYE2*) (Fig. 6C). This module may gather conserved elements of the response to darkness. Finally, networks of genes neighbouring expression for CCGs and known transcriptional regulators of the PAO/phyllobilin pathway (as in Fig. 4) were extracted and their respective position in the networks visualised (Fig. 6C, for the sake of clarity, only the three most correlated genes are shown here). Surprisingly, not all CCGs were co-expressed in a unique cluster, and not necessarily with the predicted pathway, they were shown to interact with (Fig. 6C). For example, *MYC2/3/4* and *JAZ* genes were scattered across various modules, whereas genes involved in ethylene signalling (*EIN2*, *EIN3* and *ERF17*) were mostly grouped within the blue module. As shown before (Hickman et al., 2017), differences in the networks of JA-related genes may be explained by the interplay between several factors that are linked to the treatment and genotypes used here and that are thus represented in these data, i.e. dark treatment, pheide *a* and JA.

**Figure 6.**
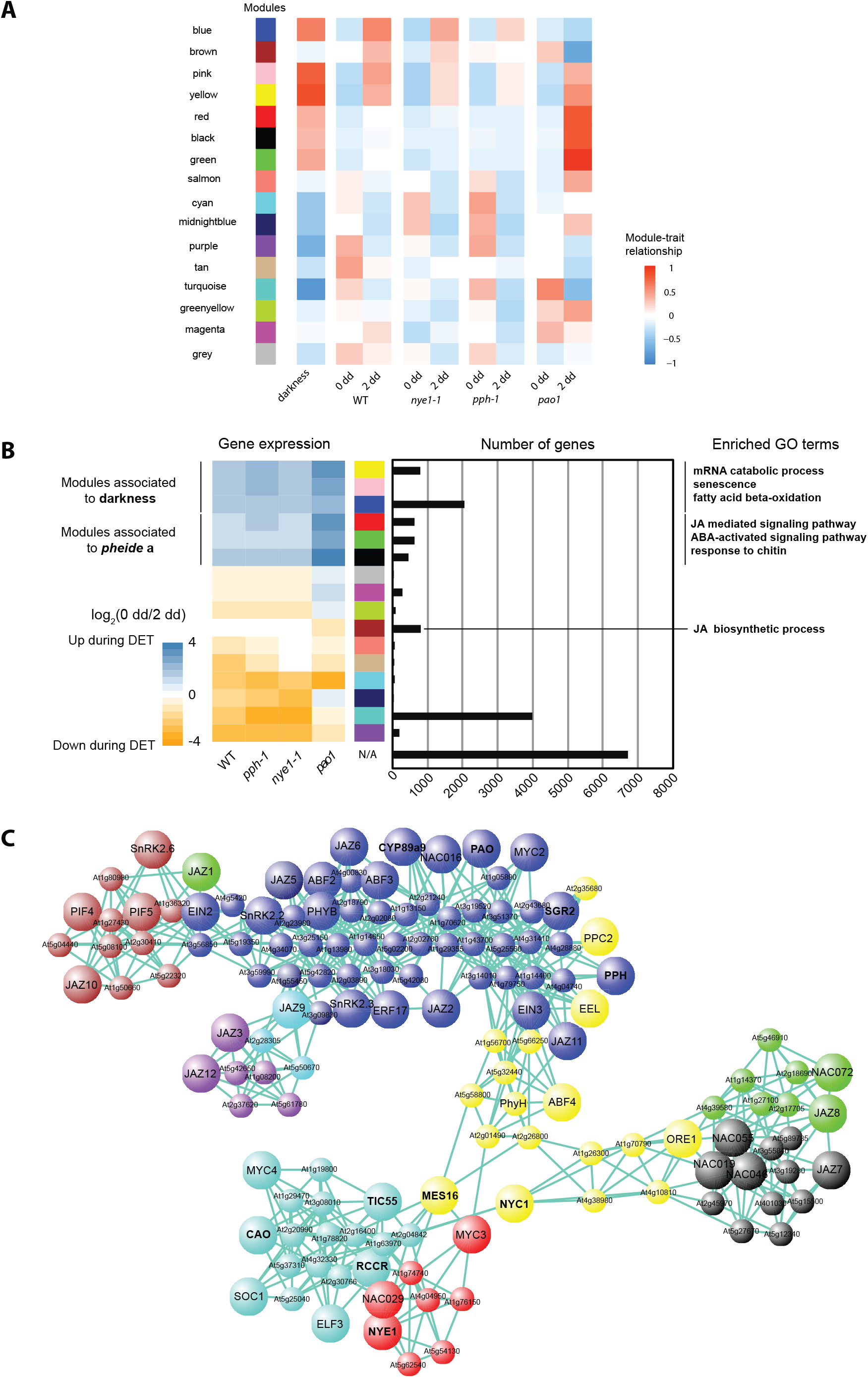
Weighted gene co-expression analysis (WGCNA) sheds new light on the regulation of the PAO/phyllobilin pathway. **A** Heat map showing a module-sample association matrix. Each row corresponds to a module. The heat map colour code from blue to red indicates the correlation coefficient between the module and either the treatment (first column; darkness) or the genetic background. **B** Patterns of expression (left panel) and size (right panel) of gene co-expression modules. On the left panel, heat maps indicate mean expression [log_2_ (fold change)] of the 10% most representative genes (highest connectivity) for each WGCNA module during dark-induced senescence. **C** The regulatory network of the PAO/phyllobilin pathway as exported from WGCNA and visualized in VisANT 5.51 (Hu et al., 2007). Larger nodes show the input genes (CCGs, transcriptional regulators according to Fig. 4), smaller nodes were limited to the top 3 most connected genes for each input gene, the edges represent connections between the genes. Node colours represent the module in which the genes clustered during WGCNA analysis.

Validation of the clustering approach can be seen, for example, by the fact that *NAC019*, *NAC055* and *NAC072*, clustering closely together, have already been shown to be homologs (Zheng et al., 2012). Similarly, ORE1 and ANAC046 are closely related, but act in distinct clusters, suggesting a distinct regulation mechanism as shown recently (Park et al., 2018). The WGCNA approach can also be a fruitful approach to identify new candidates in the PAO/phyllobilin pathway, like for example phytanoyl CoA 2-hydrolase (*phyH*) that was suggested to be involved in phytol chain degradation (Araújo *et al.*, 2011) and clustered within the yellow module with *MES16*and *NYC1*. Taken together, the co-expression data suggest that the PAO/phyllobilin pathway is regulated by multiple layers of transcriptional factors. This approach may help deciphering multiple gene networks involved in the regulation of chl degradation that are tightly associated with developmental cues, nitrogen levels, and biotic and abiotic stresses. Further work is necessary to confirm the relative influence of each of these clusters.

## DISCUSSION

We have shown that the PAO/phyllobilin pathway is mostly regulated at the transcriptional level during dark-induced senescence. A tight control of the expression of genes involved in this pathway is necessary to prevent possible oxidative damage due to a release of toxic tetrapyrrole breakdown intermediates. By genetically modulating the homeostasis of chl catabolites, we unravelled a retrograde signalling function for the PAO/phyllobilin pathway that uses JA-signalling to coordinate chloroplasts and the nucleus during dark-induced senescence.

### Pheide *a* Is a Key Signalling Molecule of Chloroplast Function

The *pao1* mutant has originally been identified in a screen for lines that show abnormal response to pathogens by accelerated severe cell death (Greenberg and Ausubel, 1993). The basis of this light-dependent cell death phenotype is relatively well understood (Yang et al., 2004; Pružinská et al., 2005). Pheide *a* phototoxicity is even observable in mammalian systems (Jonker et al., 2002). However, another peculiar feature of *pao1* is a light-independent cell death phenotype, whose underlying molecular basis is still unclear (Hirashima et al., 2009) (Fig. 1C). Two hypotheses were proposed to explain cell death caused by pheide *a* accumulation in the dark: it may act directly on chloroplast membrane integrity (via lipid peroxidation or increased oxidative stress levels) or it may itself be a signalling molecule regulating cell death. The *acd2-2* mutant that is deficient in the next committed step of the PAO/phyllobilin pathway, *i.e*. red chlorophyll catabolite reductase (RCCR) accumulates red chlorophyll catabolite (RCC), a linear tetrapyrrole. Surprisingly, although RCC is phototoxic like pheide a, *acd2-2* is exclusively affected in a light-dependent manner (Supplemental Fig. S3) (Greenberg et al., 1994; Pružinská et al., 2007). The main difference is that pheide a, unlike RCC, is likely trapped within the chloroplast. Indeed, miss-targeting of the cytosolic phyllobilin-modifying enzyme MES16 into the chloroplast in a *pao1* background revealed that *in vivo* pheide *a* is not a substrate for MES16. It can, therefore, be concluded that pheide *a* is unlikely to be released from the chloroplast (Christ et al., 2012), in contrast to RCC, which has been shown to (partially) localize to the vacuole (Pružinská et al., 2007).

Independent of the exact molecular basis underlying light-independent cell death in *pao1*, pheide *a* appears to have two specific properties: it is a metabolic “bottleneck” of degradation, *i.e*. once formed, chl molecules must be irreversibly degraded further, and it exhibits certain light-independent bioactive properties that act on chloroplast homeostasis. These two features render pheide *a* a very good candidate compound for sensing the rate at which chl is degraded, not only in the context of (natural/induced) senescence, but also during the pathogen-induced hypersensitive response (Mur et al., 2010). Sensing the rate at which chl is degraded is essential to coordinate various senescence processes such as nitrogen remobilisation (Hörtensteiner and Feller, 2002). Taking advantage of our large dataset, we propose a model of how pheide a-dependent signalling possibly works.

### Pheide *a* Metabolism Underpins a Specific Jasmonic Acid Response

Absence of PAO during dark incubation and the concomitant accumulation of pheide *a* seem to be characterised by enhanced gene expression of most of the genes from JA synthesis and signalling pathways, as well as by an increase in JA and many of its derivatives. One common feature among the CCG mutant lines studied here is the variation of levels/flux of pheide a: lower amounts in *nye1-1/pph-1* (formation blocked by up-stream mutations) vs. higher amounts (further degradation blocked) in *pao1*. We propose a model, in which the quantity of pheide *a* that accumulates in/flows through the PAO/phyllobilin pathway at a defined time may act as a signal that triggers a specific JA response (Fig. 7).

**Figure 7.**
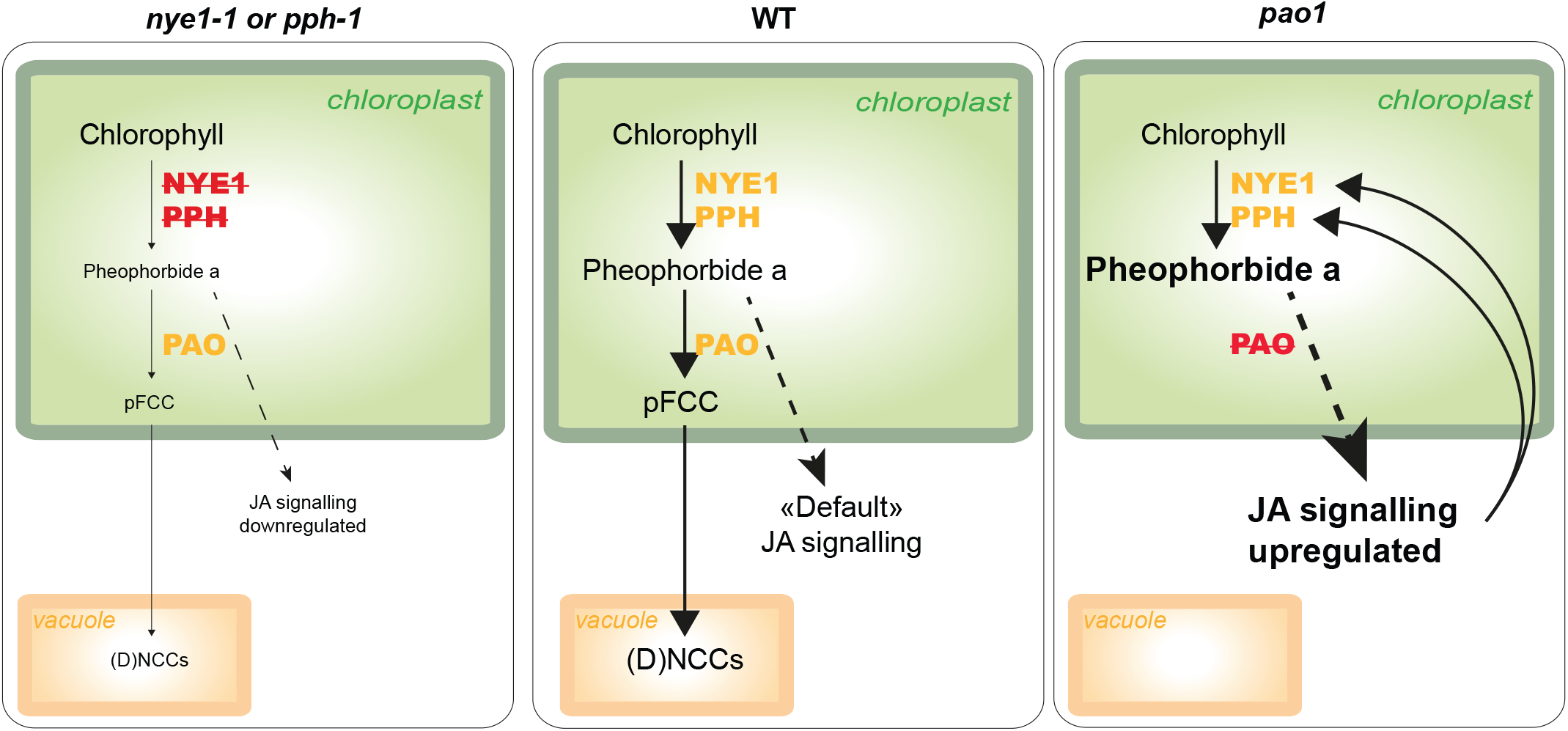
Model illustrating the influence of pheophorbide *a* homeostasis on JA signalling. The middle panel shows the PAO/phyllobilin pathway under normal senescence conditions, leading to the complete degradation of chlorophyll to vacuole-localized phyllobilins. Left and right panels show modulation of catabolite homeostasis caused by mutations of either *nye1-1* or *pph1* (left panel) or *pao1* (right panel), and the respective observed downstream modulation of the JA response (hatched arrows). Arrow sizes schematically represent relative flux (metabolite) and response (JA signalling) intensities. Among the few genes differentially expressed in *nye1-1* and *pph-1*, JAZ genes were downregulated compared to WT. On the other hand, in *pao1*, JA biosynthesis and signalling genes as well as some JA bioactive derivatives were induced.

Noteworthy, JA levels increase during dark-induced senescence in WT plants, but this is not necessarily followed by a coordinated JA response (Seltmann et al., 2010b). Prolonged darkness treatment is thought to induce degradation of chloroplast membranes and leads to an increase in lipid β-oxidation (Seltmann et al., 2010a). Breakdown of membrane lipids leads to a considerable increase in free *α*-linolenic acid, a precursor of JA. However, in *pao1*, even if the extent remains unknown to which dark-dependent pheide *a* accumulation may damage chloroplast membranes, we could observe a massive increase of JA, way above the levels of WT, and a change in most genes associated with the JA response. All these elements are strongly indicative of a fully coordinated and transduced JA response. This response is partially similar to JA responses observed during defense processes against insects, necrotrophic pathogens or ozone stress (Howe et al., 2018). It is important to note, however, that increase in *LOX* expression resulting in increased LOX activity might in itself be sufficient to increase lipid peroxidation and changes in α-linolenic acid availability and modulate oxidative stress that in turn would deteriorate chloroplast homeostasis (Wasternack, 2014; Mata-Perez et al., 2015; Wasternack and Feussner, 2018).

The pheide *a*-dependent JA response could also be differentiated across the CCG mutants studied here depending on their level of impairment in the pathway. Indeed, expression of several *JAZ* genes was reversed in *pao1* as compared to *nye1-1/pph-1* (Fig. 4). Simultaneous increase in *JAZ* expression and JA accumulation might be indicative of a heavy transduction load of the pathway (*i.e*. JAZ degradation by SCF^COI1^) making new synthesis of JAZ transcriptional inhibitors necessary.

The co-expression patterns of the JA signalling networks confirm that some of the JA signalling elements were co-expressed with various CCGs, but also illustrate the underlying complexity of the JA response (Fig. 6) (Hickman et al., 2017). For example, not all MYC2 target genes are triggered in the same manner after pheide *a* accumulation: the defensin gene *PDF1.2* (At5g44420), a known marker of ET and JA (Lorrain et al., 2003), or *VSP1* (At5g24780), induced by wounding and JA, were not differentially expressed in any of the CCG mutants, whereas *PR4* (At3g04720), a pathogenesis-related gene, was significantly overexpressed during dark incubation in *pao1* exclusively. Further studies deciphering the various interacting sub-networks involved in the JA response upon developmental and various pathogenic cues will be needed to possibly explain these discrepancies.

Several independent pieces of evidence point towards an effect of chl degradation on JA signalling. For example, *NYE* mutants are less sensitive to pathogens and have a reduced JA response compared to WT (Mecey et al., 2011). Thus, preventing chl catabolites to be metabolised by the pathway by mutating *NYE* can apparently have some protective effect. Interestingly, *pxa1*, a mutant impaired in a peroxisomal ABC transporter essential for fatty acid degradation, accumulates α-linolenic acid and pheide *a* during extended darkness (Kunz et al., 2009; Nyathi et al., 2010). This further supports the idea of an intricate link between chloroplast membrane integrity, levels of pheide *a* and JA signalling.

### Pheide *a* Signalling: Another Porphyrin-Based Retrograde Signal?

Tetrapyrrole intermediates, like Mg-protoprophyrin IX or heme have been long suggested to be involved in plastid-to-nucleus retrograde signalling (Chi et al., 2013). Coordination of chloroplast function with nuclear genome expression is equally important during early developmental stages as during senescence. Interestingly, *hy1-101* (also referred to as *gun2*), a mutant deficient in HEME OXYGENASE 1 (HO1) that catalyses heme degradation into biliverdin IX as a key step for phytochrome chromophore biosynthesis, constitutively accumulates high amounts of JA and high levels of JA-responsive genes (Zhai et al., 2007). Deficiency in HO1 leads to the accumulation in the chloroplast of protoporphyrin IX, a circular porphyrin with a structure similar to pheide a. It remains to be shown, to which extent this phenotype is similar to the one observed in *pao1* and whether porphyrin-induced JA responses could be effectively coordinated signalling mechanisms, by which the status of the chloroplast can be further transduced to the nucleus during both synthesis and degradation of chl (Lin et al., 2016).

## CONCLUSION

Taken together, our data show that the homeostasis of chl derivatives in the PAO/phyllobilin pathway impacts leaf metabolism; specifically, the rate of accumulation of pheide *a* triggers JA-related responses that, to a certain extent, mimic pathogen responses. The JA-induced transcription factor MYC2 is involved in PAO/phyllobilin pathway activation by directly binding to the promoter of various CCG, like *PAO*, *NYC1* and *NYE1* (Zhu et al., 2015; Kuai et al., 2018). Here, we show a positive feedback loop mediated by pheide a, that in turn activates JA-responsive genes. While JA signalling is central to senescence regulation, this suggests an additional signalling function of the PAO/phyllobilin pathway besides default porphyrin detoxification.

Pheide *a* has likely been recruited during evolution as a signalling molecule of the chloroplast metabolic status, due to its particular position within the chl degradation pathway and because of its intricate chloroplast toxicity. Pheide *a* signalling may act via accumulation of JA and its bioactive derivatives that in turn induce JA-dependent responses. However, the exact molecular mechanism, in particular the nature of the retrograde signal(s) that links chloroplast pheide *a*-sensing to nuclear variation in gene expression remains to be identified. To the best of our knowledge, this report is the first to postulate retrograde signalling during leaf senescence. We show how critical the control of such signals is during late leaf development stages. This proposed mechanism allows a chloroplast-controlled remodelling of the nuclear transcriptome and aims at an efficient coordination of the cellular fate during senescence.

## MATERIALS AND METHODS

### Plant Material

WT and CCG mutant lines, *i.e*. the T-DNA lines *pao1* (Pružinská et al., 2005) and *pph-1* (Schelbert et al., 2009) and the EMS line *nye1-1* (Ren et al., 2007), were grown in short day condition (8 h light/16 h dark, 23°C, 65% humidity) for eight weeks. At least four leaves n°8 for each triplicates were harvested and frozen in liquid nitrogen at 0 days in the dark (dd) and after 2 days incubation on H_2_0-soaked filter paper in complete darkness at 23°C.

### RNA Isolation and Sequencing

RNA was isolated using RNAeasy minikit (Qiagen) together with on-column DNAse treatment. Quality was assessed using Bioanalyzer RNA nanochip (Agilent). Three replicates samples for each condition were multiplexed randomly on two lanes (12 samples per lane) of HiSeq 2500 (Illumina).

### Read Processing and Gene Expression Analysis

Single-end 100 bp reads were subjected to adapter trimming and removal of low quality bases in leading, trailing and sliding window (4 bp) mode with Trimmomatic v0.35 (Bolger et al., 2014). Reads shorter than 40 bp after trimming were discarded. Remaining reads were aligned to the protein-coding transcripts from the ENSEMBL release of the TAIR10 *Arabidopsis thaliana* transcriptome (Swarbreck et al., 2008) using Bowtie v1.0.1 (Langmead, 2010). Expression of genes and transcripts was quantified using RSEM v1.2.11 taking into account strand-specific information (Li and Dewey, 2011). Differential expression was estimated using EBSeq by estimating the posterior probability of genes to be differentially expressed across all conditions (Leng et al., 2013). Coverage data were visualized using IGV viewer 2.3.34 (Thorvaldsdottir et al., 2013) using RSEM-generated .bam files (see Supplemental Data). Gene ontology enrichment was performed using a corrected Benjamini-Hochberg enrichment score implemented in Pageman (Usadel et al., 2006).

### Co-Expression Network Analysis

WGCNA was used to identify modules gathering genes showing similar pattern of expression across all conditions (Langfelder and Horvath, 2008). Genes below 50 mean read count were excluded, leaving 14,691 genes in the analysis. An unsigned network was constructed from a signed topological overlap matrix and module detection was performed using the default deepSplit setting of 2. In order to visualize the direct subset of genes co-regulated with CCG and selected regulatory gene candidates, subnetworks were generated and visualized using VisANT 5.51 (Hu et al., 2007). In order to evaluate the extent to which expression of genes involved in the regulation of the PAO/phyllobilin pathway (all present in Fig. 4) were linked to CCE genes, subnetworks containing either of these genes (CCGs and regulators) were extracted from the WGCNA networks and the three most connected genes for each gene were displayed (Fig. 6C). Larger nodes show the input genes and smaller nodes the top three connected genes for each input gene. Edges represent connection between the genes and node colors represent the modules in which the genes clustered.

### Chlorophyll Extraction

Chl was extracted from liquid-nitrogen homogenised tissue using extraction buffer (90% cold acetone and 10% 0. 2 M Tris-HCl, pH 8) (Guyer et al., 2014). Chl content was determined by photospectrometry at A_649_ and A_665_. Chl concentrations were calculated as published (Strain *et al.*, 1971).

### Chlorophyll Catabolites Profiling

Metabolite profiling was performed by liquid chromatography (LC)-tandem mass spectrometry (MS) (LC-MS/MS) according to a published protocol (Christ et al., 2016). Briefly, leaf samples from 5 replicates were harvested, frozen and homogenized in liquid nitrogen. Metabolites were extracted in five volumes of ice-cold extraction buffer [80% methanol, 20% water, 0.1% formic acid (v/v/v)] and centrifuged (5 min at 14,000 rpm, 4°C). Supernatants were then analyzed by LC-MS/MS.

Samples were run on an Ultimate 3000 Rapid Separation LC system (Thermo Fisher Scientific) coupled to a Bruker Compact ESI-Q-TOF (Bruker Daltonics). The system consisted of a 150 mm C18 column (ACQUITY UPLC BEH, 1.7 μm; Waters Corp., Milford, MA, USA). In order to efficiently separate phyllobilins, the following gradient of solvent B [acetonitrile with 0.1% (v/v) formic acid] in solvent A [water with 0.1% (v/v) formic acid] was run at a flow rate of 0.3 mL min^-1^: 5% B for 0.5 min, 5% B to 100%B in 11.5 min, 100% B for 4 min, 100% B to 5% B in 1 min and 5% B for 1 min. Pheide *a* and phyllobilins were quantified from extracted ion chromatograms as relative peak areas using QuantAnalysis (Bruker Daltonics).

### Determination of Phytohormones

Extraction was performed as previously described for lipids (Matyash et al., 2008) with some modifications. Five replicates were used for each condition and each time point. Plant material (100 mg) was extracted with 0.75 mL of methanol containing 10 ng D5-JA (C/D/N Isotopes Inc., Pointe-Claire, Canada), 30 ng D5-oPDA, 10 ng D4-JA-Leu (both kindly provided by Dr. Otto Miersch, Halle, Germany) as internal standards. After vortexing, 2.5 mL of methyl-tert-butyl ether (MTBE) were added and the extract was shaken for 1 h at room temperature. For phase separation, 0.6 mL H2O were added. The mixture was incubated for 10 min at room temperature and centrifuged at 450 g for 15 min. The upper phase was collected and the lower phase re-extracted with 0.7 mL methanol/water (3:2.5, v/v) and 1.3 mL MTBE as described above. The combined upper phases were dried under streaming nitrogen and re-suspended in 100 μL of acetonitrile/water (1:4, v/v) containing 0.3 mM NH_4_COOH (adjusted to pH 3.5 with formic acid).

Reversed phase separation of constituents was achieved by LC using an ACQUITY UPLC system (Waters) equipped with an ACQUITY UPLC HSS T3 column (100 mm × 1 mm, 1.8 μm; Waters). Aliquots of 10 μL were injected. Elution was adapted from a published procedure (Balcke et al., 2012). Solvent A and B were water and acetonitrile/water (9:1, v/v), respectively, both containing 0.3 mM NH_4_COOH (adjusted to pH 3.5 with formic acid). The flow rate was 0.16 mL min^-1^ and the separation temperature held at 40°C. Elution was performed with two different binary gradients. Elution profile 1 was as follows: 10% B for 0.5 min, to 40% B in 1.5 min, 40% B for 2 min, to 95% B in 1 min, 95% B for 2.5 min; elution profile 2: 10% B for 0.5 min, to 95% B in 5 min, 95% B for 2.5 min. In both elution profiles, the column was re-equilibrated in 10% B in 3 min.

Nano-electrospray ionization (nanoESI) analysis was achieved using a chip ion source (TriVersa Nanomate; Advion BioSciences, Ithaca, NY, USA). For stable nanoESI, 70 μL min^-1^ of 2-propanol/acetonitrile/water (7:2:1, v/v/v) containing 0.3 mM NH_4_COOH (adjusted to pH 3.5 with formic acid) delivered by a Pharmacia 2248 HPLC pump (GE Healthcare, Munich, Germany) were added just after the column via a mixing tee valve. By using another post column splitter, 502 nL min^-1^ of the eluent were directed to the nanoESI chip with 5 μm internal diameter nozzles. Jasmonates were ionized in negative mode at -1.7 kV (after UPLC separation with elution profile 1) and in positive mode at 1.3 kV (after UPLC separation with elution profile 2), respectively, and determined in scheduled multiple reaction monitoring mode with an AB Sciex 4000 QTRAP tandem mass spectrometer (AB Sciex, Framingham, MA, USA). Mass transitions were as previously described (Iven *et al.*, 2012), with some modifications as follows: 214/62 [declustering potential (DP) 35 V, entrance potential (EP) 8.5 V, collision energy (CE) 24 V] for D5-JA, 209/59 (DP 30 V, EP 4.5 V, CE 24 V) for JA, 237/165 (DP 45 V, EP 6 V, CE 24 V) for OPC4, 265/221 (DP 50 V, EP 6 V, CE 24 V) for OPC6, 305/97 (DP 30 V, EP 4 V, CE 32 V) for 12HSO4-JA, 338/130 (DP 45 V, EP 10 V, CE 30 V) for 12OH-JA-IIe, 352/130 (DP 45 V, EP 10 V, CE 30 V) for 12COOH-JA-IIe, 387/59 (DP 85 V, EP 9 V, CE 52 V) for 12O-Glc-JA, 325/133 (DP 65 V, EP 4 V, CE 30 V) for D4-JA-Leu, 308/116 (DP 45 V, EP 5 V, CE 28 V) for JA-Val, 322/130 (DP 45 V, EP 5 V, CE 28 V) for JA-Ile, 296/170.2 (DP 65 V, EP 4 V, CE 28 V) for D5-OPDA, 263/165 (DP 40 V, EP 5 V, CE 20 V) for dnOPDA and 291/165 (DP 50 V, EP 5 V, CE 26 V) for 12-OPDA. The mass analyzers were adjusted to a resolution of 0.7 amu full width at half-height. The ion source temperature was 40°C, and the curtain gas was set at 10 (given in arbitrary units). Quantification was carried out using a calibration curve of intensity (m/z) ratios of [unlabeled]/[deuterium-labeled] vs. molar amounts of unlabeled (0.3-1000 pmol) compound. Due to the lack of standards, only relative amounts of 12HSO4-JA, 12OH-JA-Ile, 12COOH-JA-Ile and 12O-Glc-JA were determined.

### Ion Leakage Measurements

For determining cell death in the lines during senescence, leaf discs (0.4 cm diameter) were punched with a cork-borer under green safe light, avoiding the mid vein. They were placed in a multi-well plate ion conductivity meter (Reid & Associates, South Africa) (1.5 mL H2O and two discs per well) and relative ion leakage (displayed as μS) was determined in the dark.

### Accession Number

The raw sequencing data from RNAseq are available in the ArrayExpress database (www.ebi.ac.uk/arrayexpress) under accession number (E-MTAB-6965).

## Supporting information

## List of author contributions

S.A. and S.H. conceived the original research plans; S.A. and S.O. performed most of the experiments; K.Z. and I.F. analyzed jasmonic acid metabolites; S.A. and N.F. analyzed the data; S.A. and S.H. wrote the article with contribution of all the authors.

## Supplemental Data

The following supplemental materials are available.

**Supplemental Figure S1.** Overlap between the data presented here and two independent leaf senescence transcriptome datasets.

**Supplemental Figure S2.** Dendrogram of the modules generated by WGCNA

**Supplemental Figure S3.** Electrolyte leakage data of *pao1* and *acd2-2* mutants during dark-induced senescence.

**Supplemental Figure S4.** Mapping of the RNAseq reads to genes of interest in respective mutant lines.

**Supplemental Dataset S1.** RNAseq gene expression data during DET in the four lines.

**Supplemental Dataset S2.** GO terms enrichment for each pairwise comparison of gene expression.

**Supplemental Dataset S3.** List of 50 most highly expressed genes after dark incubation.

**Supplemental Dataset S4.** Data from quantification of jasmonic acid and its derivatives used to draw Fig. 5.

**Supplemental Dataset S5.** Expression of hormone-related genes in the four lines before senescence induction.

**Supplemental Dataset S6.** GO terms enrichment of all 16 clusters originating from the WGCNA analysis and WGCNA scoring matrix for dark-treated leaves across all lines and for *pao1* after 2 dd incubation.

## ACKNOWLEDGEMENTS

We are thankful to Sirisha Aluri and Lennart Opitz from the Functional Genomics Centre Zürich for sequencing, to Otto Miersch from the University of Halle, Germany, for providing JA metabolite standards, and to Kathrin Salinger for technical assistance.

