## Supplementary material for "Pheophorbide *a*, a chlorophyll catabolite may regulate jasmonate signalling during dark-induced senescence in *Arabidopsis*"

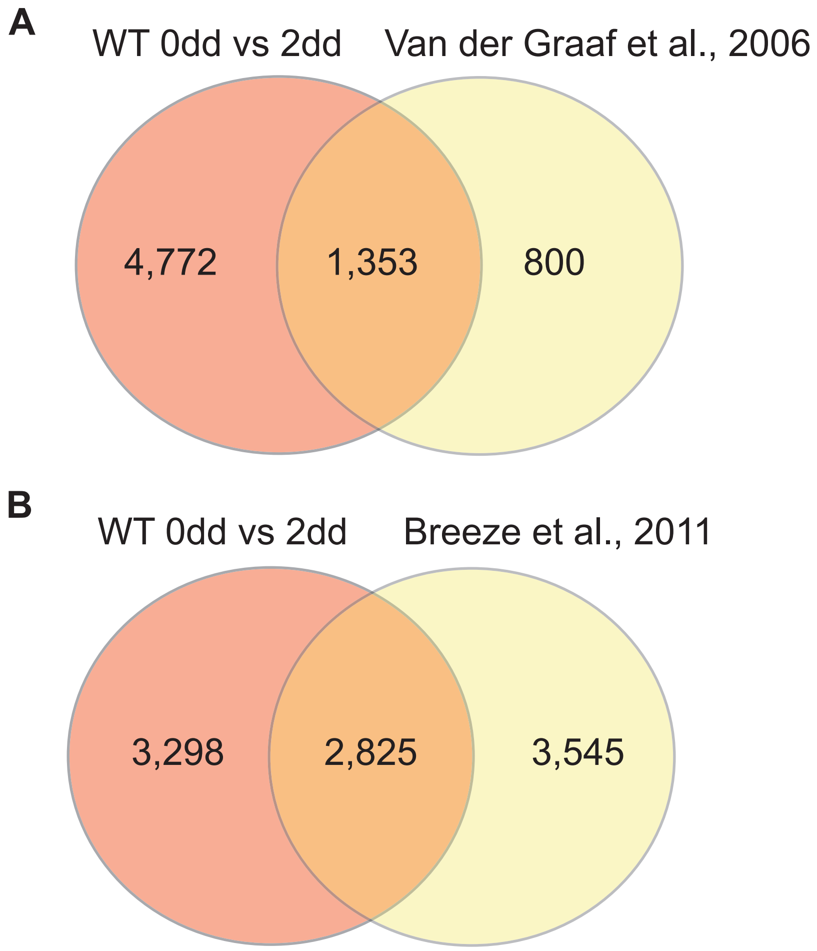


**Supplemental Figure S1. Overlap between the data presented here and two independent leaf senescence transcriptome datasets. A** Van Der Graaff et al., 2006 **B** Breeze et al., 2011.

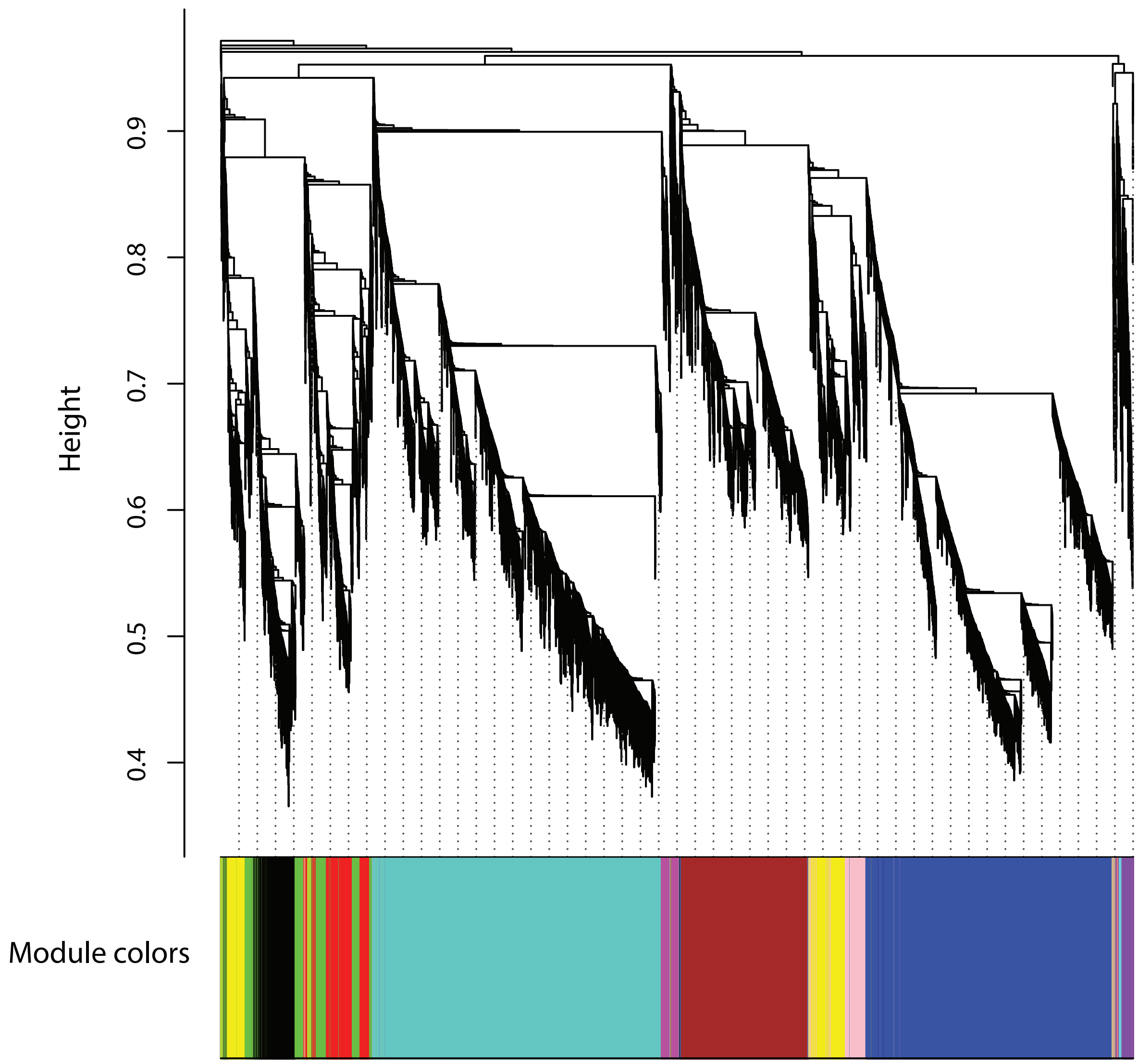


**Supplemental Figure S2. Dendrogram of the modules generated by WGCNA.**

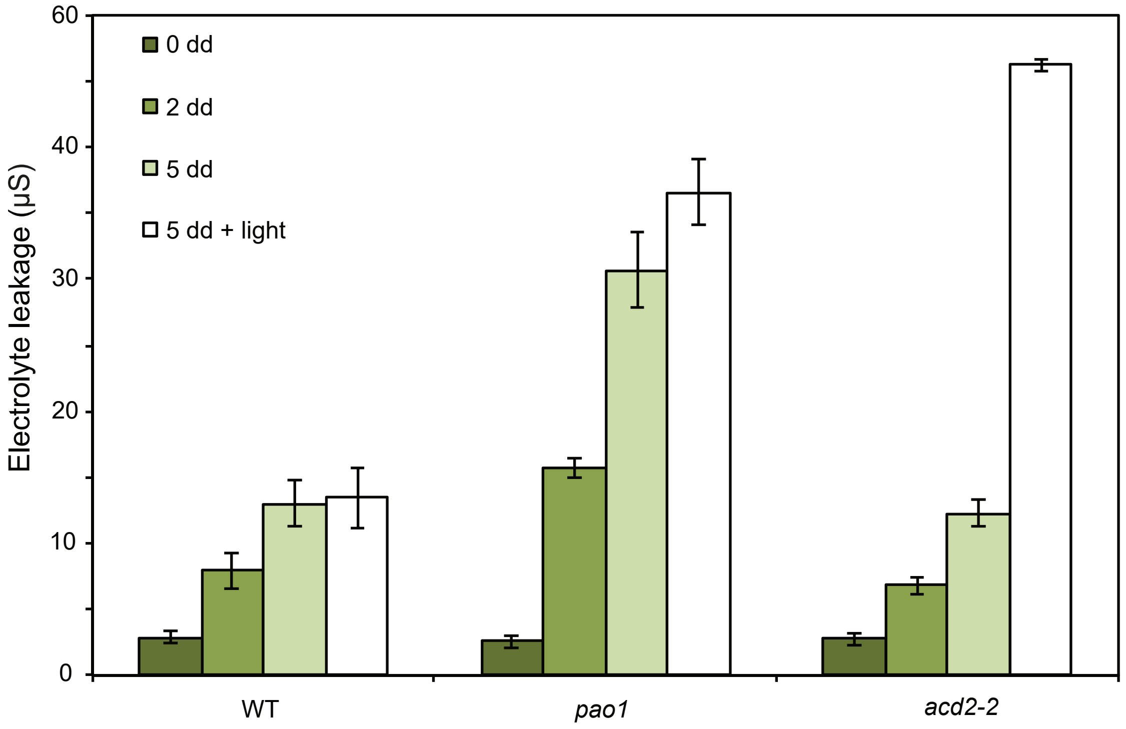


**Supplemental Figure S3. Electrolyte leakage data of *pao1* and *acd2-2* mutants during dark-induced senescence.** Data are mean values of a representative experiment with at least ten replicates. Error bars indicate SD. In this experiment, leaves were subjected to one-hour light treatment after the 5 dd incubation (white bars).

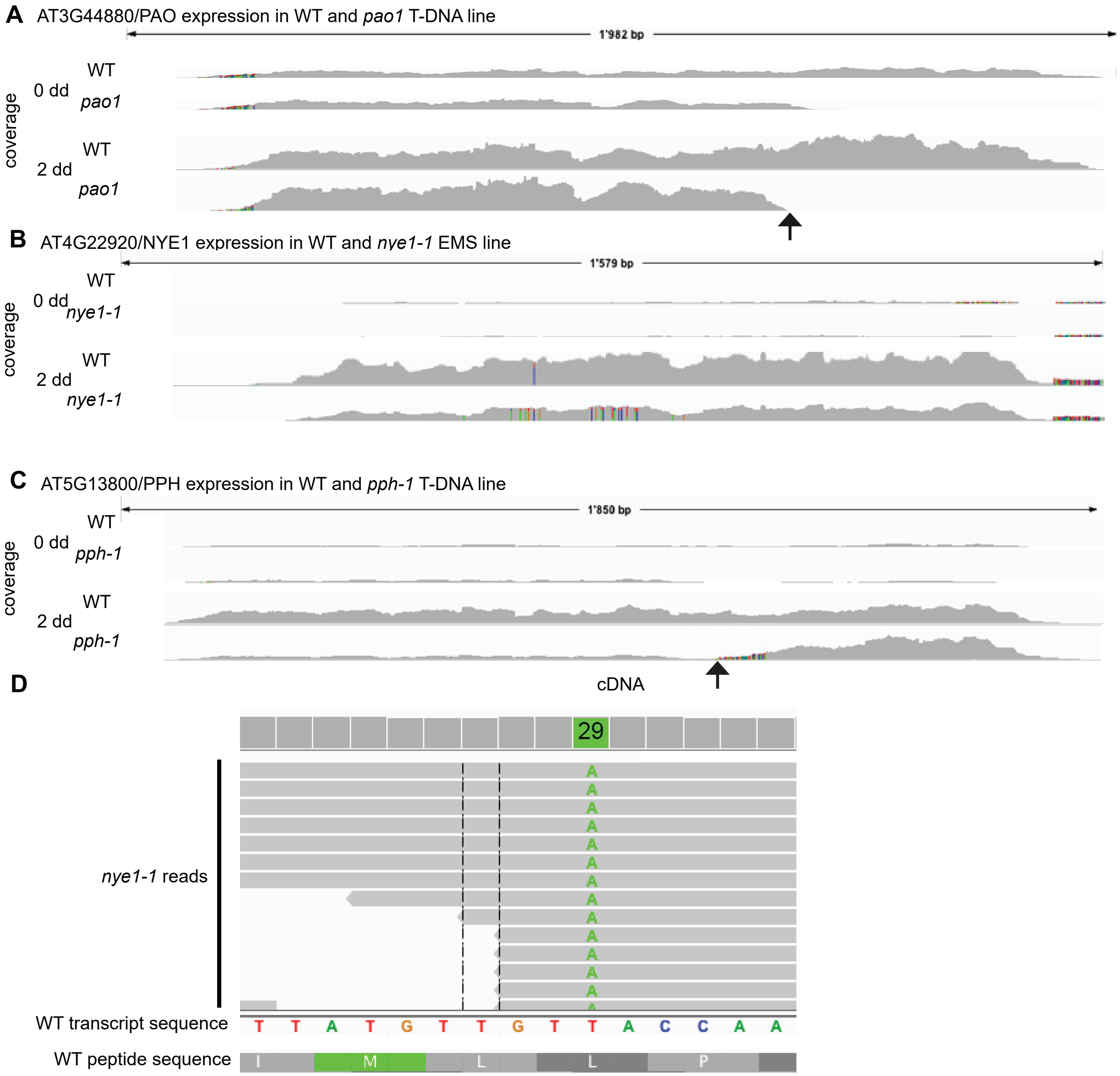


**Supplemental Figure S4. Mapping of the RNAseq reads to genes of interest in respective mutant lines.** Figures were extracted from IGV viewer. Mapping data are shown for only one replicate for each condition.
